# The causal role of β-oscillations in maintaining perception-action representations

**DOI:** 10.64898/2026.05.28.725394

**Authors:** Bernhard Pastötter, Paula M.A. Giebel, Phillipp P. Schummers, Christian Frings, Florian H. Kasten

**Author notes:** Corresponding author: Florian H. Kasten.

## Abstract

The binding of perception and action features into common representations, so-called event files, is a core mechanism supporting goal-directed behavior. Recent work using electroencephalography has presented correlational evidence linking the maintenance of perception-action representations to oscillatory activity in the β-frequency range. Here, we used transcranial alternating current stimulation (tACS) to test whether β-oscillations play a causal role in event-file maintenance. Participants performed a distractor-response binding task with a sequential prime-probe structure while receiving either active β-tACS over the left occipito-parietal cortex or a novel shunt stimulation designed to produce comparable sensory effects but substantially weaker cortical stimulation. Prime-probe intervals were varied to manipulate event-file maintenance. β-tACS did not modulate behavioral binding effects during stimulation. However, we observed a significant aftereffect on behavioral distractor-response binding after stimulation. This effect was restricted to short prime-probe intervals, where event files are typically still available, and was absent at longer intervals, where event files commonly decay. These findings provide causal evidence that β-oscillations contribute to the stabilization of perception-action representations over time. More broadly, they support the view that β-oscillations help maintain the current cognitive-motor state and preserve the status-quo in action control.

## 1 Introduction

The integration of perception and action is a key ability for goal-directed behavior. Prominent frameworks from cognitive psychology propose that perception, motor programs, and sensory effects of actions are bound into a common representation. This representation is conceptualized as an event file [1–4]. In this event file, features of an object such as its color or shape are represented together with actions performed on or with the object (e.g., manual responses, rotations, grasping, etc.). If any feature of the object or action is reencountered, the entire event file is retrieved, leading to distinct performance benefits or costs, depending on whether the same or a different action is required from the first to the second encounter [5,6].

The binding and retrieval in action control (BRAC) framework is a process model aiming to describe the processes of binding perception and action into event files, maintaining them as memory traces, and retrieving them upon repetition of a stimulus or response feature [2]. Recent years have seen an increased interest in the neural correlates underlying perception-action integration [7]. It has been proposed that theta and gamma oscillations might be implicated in the binding and retrieval of event files [7–9]. Alpha oscillations may affect binding and retrieval indirectly via attentional modulation. After binding, event files are subject to decay over time, remaining available for only a few hundred milliseconds to seconds [10]. However, the exact time-course of the decay remains unclear and is possibly subject to large inter-individual differences [11]. Recent evidence from electroencephalography (EEG) indicates that the maintenance of event files is linked to β-oscillations in occipito-parietal [12] and frontal cortices [13]. Stronger β-band synchronization following actions has been shown to be associated with stronger and longer-lasting maintenance. Dysfunctional β-oscillations seem to be related to hyper-binding in functional movement disorders [13].

EEG can provide evidence for the involvement of β-oscillations in event-file maintenance and may indicate their neural sources. This evidence is, however, correlational in nature and does not allow conclusions if β-oscillations play a causal role in maintaining perception-action representation or merely reflect epiphenomena of the underlying neural computations [14,15]. In order to resolve such causal involvement, it is necessary to actively manipulate β-oscillations and test if behavior changes [14]. Transcranial alternating current stimulation (tACS) is widely used in neuroscience research to manipulate endogenous brain oscillations to study their role in cognition [16]. The technique works via the application of weak alternating currents through two or more electrodes placed on the scalp. It makes use of the principles of neural entrainment to manipulate endogenous brain oscillations in a frequency-specific manner [17,18]. Besides effects occurring during stimulation due to neural entrainment, tACS is capable of inducing effects that outlast stimulation by several minutes or even up to over an hour [19,20]. These aftereffects have been linked to processes of spike timing dependent plasticity [21–23], are behaviorally relevant, and capable of changing event-related brain oscillations [24,25].

The current study aimed to investigate the causal role of β-oscillations in the left occipito-parietal cortex for event-file maintenance. Prior EEG evidence suggests that β-oscillatory activity in this region is particularly related to the maintenance of bindings between actions and task-irrelevant stimulus information [12], whereas β-activity in the frontal cortex has been linked more strongly to bindings involving task-relevant stimulus information [13] . To specifically assess bindings between responses and task-irrelevant distractor features, we employed the distractor-response binding (DRB) paradigm with sequential prime-probe structure [6]. Accordingly, the left parietal cortex was considered the most relevant stimulation target. We applied β-tACS or a novel inactive shunt stimulation [26,27], while participants performed the DRB task. We manipulated event file maintenance by changing the interval between prime response and the subsequent probe stimulus (Response-Stimulus-Interval, RSI; 500-ms vs. 2000-ms). We hypothesized that tACS in the β-frequency range would lead to reduced decay of DRB effects across increasing RSIs compared to shunt stimulation. More specifically, we expected a significant effect of stimulation on behavioral binding, reflected in larger DRB effects during β-tACS than during shunt stimulation, especially during the stimulation period (online). Given prior evidence for tACS aftereffects, this enhancement may also persist beyond the stimulation phase and thus be detectable in the post-stimulation period (offline) and may be affected by the RSI because behavioral DRB effects are typically observed for a shorter RSI (500-ms) but not for a longer RSI (2000-ms; e.g. [6,12], but see [11]). Such a pattern would signify a distinct effect of β-tACS in event-file maintenance rather than a general effect on performance or event-file binding or retrieval.

## 2 Methods

### 2.1 Participants

Forty-four participants (28 identified as female, mean age = 23.9 ± 4.0 years) were recruited for the study. Sample size was calculated based on an a-priori power analysis using G*Power aiming for a power (1-β) of .95 and an α-level of .05 (one-sided). Participants were screened for contraindications for transcranial electrical stimulation (Antal et al., 2025), including metal implants in the head, history of neurological or psychiatric disorders, current medication, or a history of head trauma, etc. The study was approved by the local ethics committee at the University of Trier (EK No. 21/2025) and was performed in accordance with the Declaration of Helsinki. The study was preregistered on AsPredicted.org (No #249844, https://aspredicted.org/nb79m2.pdf). Deviations from the original analysis plan will be outlined throughout the manuscript.

### 2.2 Stimulus Material

Participants performed a DRB task consisting of sequential prime-probe episodes (**Fig. 1a**). In each black display, a target letter was presented centrally at fixation and flanking distractor letters were presented to the left and right of the target. Target stimuli were the letters D, F, J, and K. The letters D and F were mapped to a left-hand response, whereas the letters J and K were mapped to a right-hand response. Participants responded with the left and right index fingers using the keys C and M on a standard QWERTZ keyboard.

**Figure 1:**
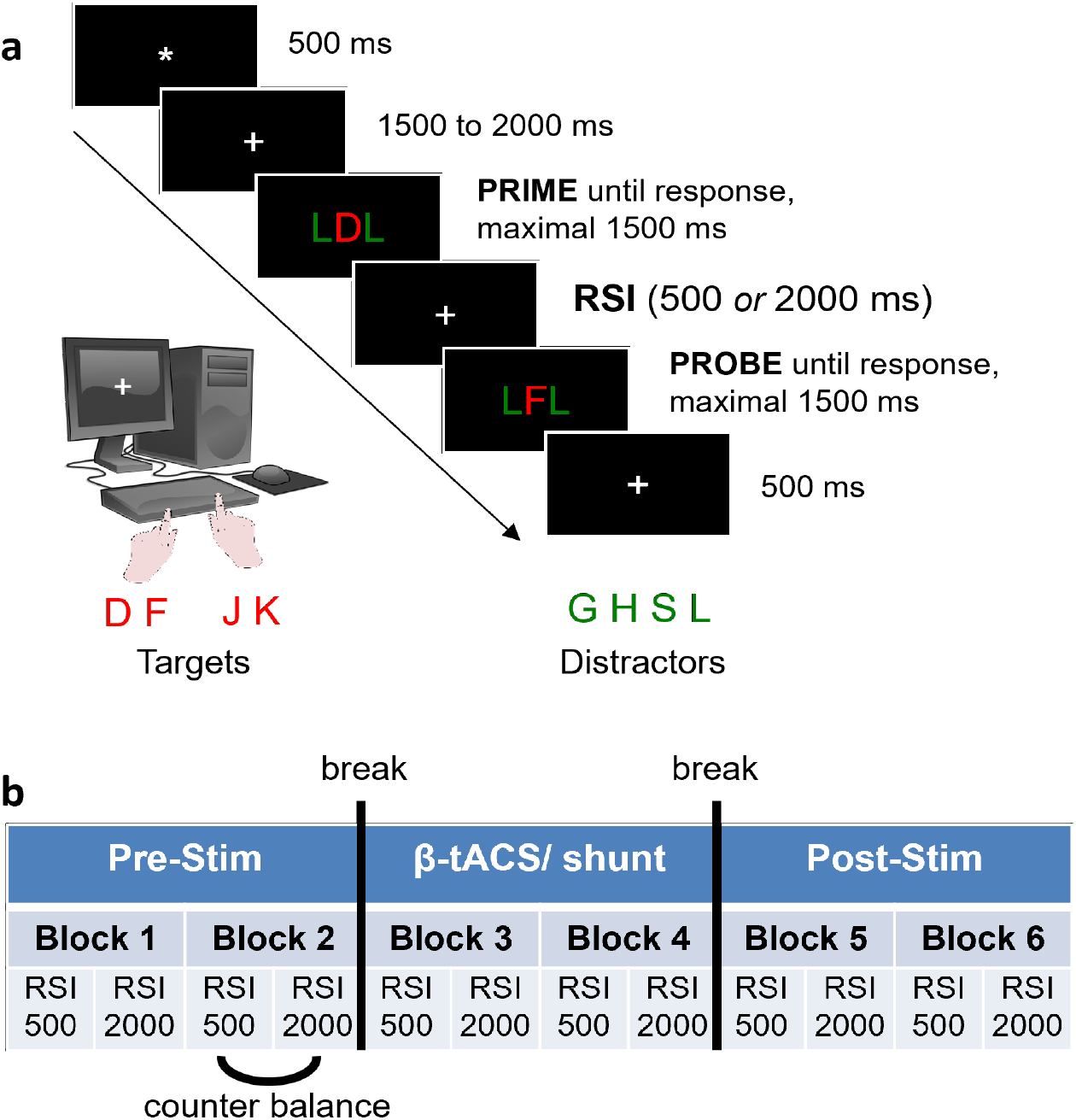
Paradigm. **(a)** Participants performed a distractor-response binding task on a computer screen. Trials consisted of a PRIME and a PROBE display. Participants were instructed to respond to the red letters in the center of the screen with their left and right index finger. Each trial consisted of a PRIME and a PROBE display presented at a fixed intervals of 500-ms or 2000-ms after PRIME response. **(b)** Time-course of the experiment. Participants performed six blocks if the task in both experimental sessions. Each block consisted of an RSI 500 and an RSI 2000 condition, presented in counterbalanced order across participants (either RSI 500 first or RSI 2000 first in each block). TACS or shunt stimulation were applied during blocks 3 & 4 of the experiment.

Different target letters were used in prime and probe displays. Target combinations were controlled to form one-to-one mappings both for response repetitions (D→F, F→D, J→K, K→J) and for response changes (D→K, K→D, F→J, J→F). Distractor stimuli were the letters G, H, S, and L. Distractors were always presented bilaterally and were never mapped to a response. Prime-probe target combinations were arranged to realize response repetitions (RR) and response changes (RC), whereas distractor identity either repeated (DR) or changed (DC) from prime to probe.

### 2.3 Experimental design

The experiment employed a within-participant design with the factors STIMULATION (active β-tACS vs. shunt control), Response Relation (RR vs. RC), Distractor Relation (DR vs. DC), and RSI (RSI: 500-ms vs. 2000-ms). Response Relation and Distractor Relation were varied orthogonally, resulting in four equally likely prime-probe conditions: RRDR, RRDC, RCDR, and RCDC.

Each participant completed two sessions on separate days, receiving active stimulation in one session and shunt stimulation in the other, with session order counterbalanced across participants. Experimental sessions were performed at least 4 days apart to avoid carry-over effects of stimulation [19,29]. Within each session, the task comprised six experimental blocks. Blocks 1 and 2 served as baseline blocks before stimulation, blocks 3 and 4 were completed during stimulation, and blocks 5 and 6 assessed aftereffects of stimulation. Each block consisted of 96 trials, including 48 trials with a 500-ms RSI and 48 trials with a 2000-ms RSI.

The block definition deviates from the preregistration, as it was originally planned to collapse blocks 1 & 2, 3 & 4 and 5 & 6 into baseline, stimulation and post-stimulation blocks, respectively. However, when inspecting the baseline data, we recognized that the DRB effect was absent in the first half of the pre-stimulation baseline (block 1) and built up in the second half (block 2, see section 3.1). As the presence of a DRB effect before stimulation is essential for its modulation with tACS, we deemed a more fine-grained separation into six blocks more appropriate for the remaining analyses. The approach further allows us to trace stimulation effects over time with a better temporal resolution.

### 2.4 Procedure

At the beginning of each session, participants were seated in front of the stimulus monitor, housed in a sound isolated, dimly lit chamber, and instructed to place their left index finger on the C key and their right index finger on the M key of the keyboard. They were told to respond to the identity of the centrally presented target letter as quickly and accurately as possible while ignoring the flanking distractors. Before the start of the experimental blocks, participants completed a short practice phase to familiarize themselves with the task. The practice phase consisted of 24 trials with equally likely prime-probe conditions.

Each trial consisted of a prime-probe sequence (**Fig. 1a**). A trial began with a central fixation display showing an asterisk for 500-ms, followed by a jittered fixation interval showing a fixation cross for 1500 to 2000-ms. Next, the prime display was presented, consisting of the target letter in the center and two identical distractor letters on the left and right. The prime display remained on the screen until the participant responded or 1500-ms had elapsed. Participants performed a short practice block before the main experiment. If the prime response was incorrect or if no response was given within 1500-ms, a 1500-ms feedback display was presented, showing the German words “Fehler!” (error!) or “Schneller!” (faster!), respectively. Thereafter, a fixation cross was shown for the duration of the current RSI condition, that is, either 500-ms or 2000-ms. Subsequently, the probe display appeared, again containing a central target and two flanking distractors. The probe display also remained on the screen until a response was made or 1500-ms had elapsed. Again, feedback was provided if responses were incorrect or too slow. Finally, a fixation cross was shown for 500-ms before the next trial began with an asterisk. While the trial structure remained identical, no feedback was provided in trials in the main experiment.

The screen background was black, whereas asterisks and fixation crosses were white. Target letters were presented in red and distractor letters in green. Target and distractor letters were approximately 0.95° in size and were displayed in bold Courier New font. Participants’ viewing distance was 60 cm.

Short breaks were provided between the six blocks of each session, including an 11 s countdown from 10 to 0. In addition, there was a longer break before and after tACS was initialized (after blocks 2 and 4). Within each block, all predefined prime-probe combinations were realized in a counterbalanced manner. Across the six blocks, each session comprised 576 experimental trials overall, with 288 trials in the 500-ms RSI condition and 288 trials in the 2000-ms RSI condition. Figure 1b provides an overview of the time course of an experimental session. Stimulus presentation and response registration were controlled using E-Prime 3.0 (Psychology Software Tools, Pittsburgh, PA, USA).

### 2.5 Electrical Stimulation

Participants received either active tACS at β-frequency (21 Hz), or a novel shunt control stimulation across the two experimental sessions. Shunt stimulation uses the same stimulation intensity, frequency and duration as active stimulation, but applies stimulation via electrodes placed too close to induce a substantial electric field in brain tissues [26]. In contrast to conventional sham stimulation, where tACS is applied for a few seconds before being switched off, shunt stimulation is applied continuously. This allows for peripheral stimulation and sensations comparable to active stimulation, thus offering better control of peripheral effects, which may partly drive brain stimulation effects [27,30]. Stimulation was applied via two surface conductive rubber electrodes connected to a neuroConn MC Stimulator (neuroCare Group GmbH, Illmenau, Germany). Electrodes were connected via a conductive paste (Ten20 conductive paste, Weaver & Co., USA). Active stimulation was applied using a small (20 mm diameter) circular electrode placed centered over the target area at position P5 of the International 10-20 system. A large concentric ring electrode with an inner diameter of 90 mm and an outer diameter of 110 mm symmetrically placed around it served as the return electrode (**Fig. 1a**). The configuration was chosen to maximize the electric field over the left parietal cortex. This region has previously been associated with β-band oscillations related to the maintenance of distractor-response bindings [12]. For the shunt montage, the outer ring electrode was replaced by a smaller ring with an inner diameter of 30 mm and an outer diameter of 48 mm (**Fig. 2b**).

**Figure 2:**
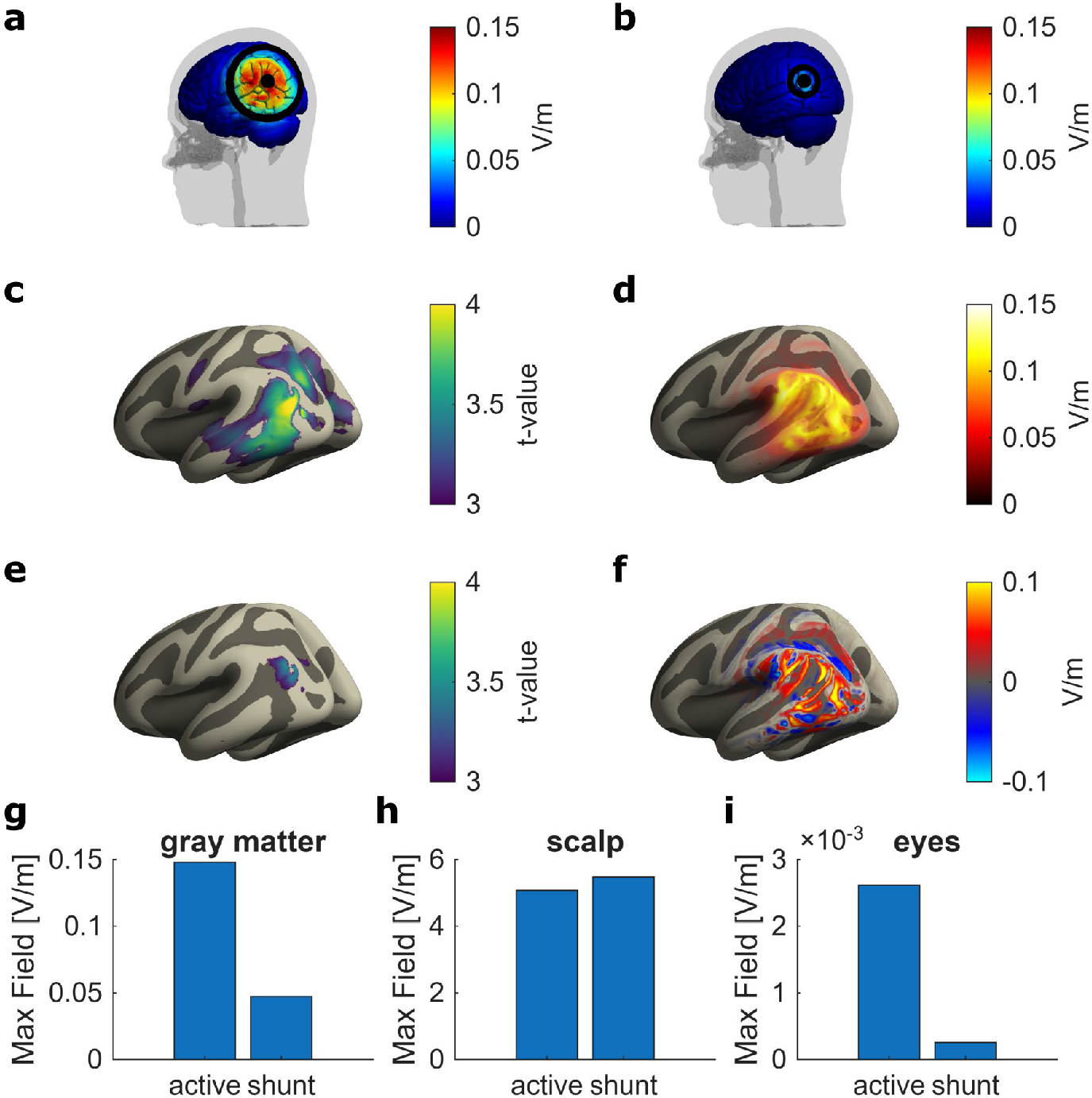
Electrode montages. **(a)** Electric field magnitude across the cortex of the MNI152 template during active stimulation centered over P5. Electrodes are depicted in back. **(b)** Electric field magnitude in the same brain model when using a smaller concentric return electrode to induce a shunting. **(c)** Source localized event-related β-source during event-file maintenance found in Pastötter et al. (2021) shown on an inflated freesurfer fsaverage brain. **(d)** Electric field magnitude shown in (a) projected onto the inflated fsaverage for better comparison with β-sources. **(e)** Reduced β-activity during longer maintenance periods reported in Pastötter et al. (2021). **(f)** Normal component of the induced electric field during active stimulation. **(g)** Comparison of peak electric fields induced to gray matter tissue by active and shunt montage. Electric fields are 3-fold smaller during shunt compared to active stimulation. **(h)** Comparison of peak electric fields induced to the scalp by both montages. Both induce comparable fields to the scalp. **(j)** Electric field induced to the eyeballs by both montages. While there is a marked difference in the induced electric field between montages, the overall field strength of both montages is very low, likely avoiding effects on retinal cells and the induction of phosphenes.

We performed electric field calculations on the MNI152 standard brain using SimNIBS 4.5 [31] to verify that our active montage maximizes the electric field over previously identified β-sources correlating with DRB (**Fig. 2a**), and to compare the exposure of the region to the electric field induced by the shunt montage (**Fig. 2b**). The induced electric field shows a strong overlap with the β-sources found in [12] (**Fig. 2 c-f**). In comparison to the active montage, our shunt montage induces an electric field to cortical tissue that is almost 3-fold weaker (**Fig. 2g**), while inducing a comparable electric field in the scalp, which ensures similar perceptual effects (**Fig. 2h**) during active stimulation and shunt. In addition, electric fields in the eyes are weak in both montages, minimizing the likelihood of inducing phos-phenes which may interfere with behavioral experiments [32]. Both simulations were run with a current injection of 2 mA (peak-to-peak). For the actual stimulation, a current injection of 2 mA (peak-to-peak) was aimed at, but intensity was reduced if participants reported discomfort due to the stimulation sensations. On average, participants were stimulated with 1.91 mA ± 0.22 mA (range: 1.1 – 2.0 mA) in the verum and 1.93 mA ± 0.22 mA (range: 1.0 – 2.0 mA) in the shunt condition.

After each session, participants were asked to fill out a questionnaire regarding common side effects of stimulation (itching, burning, tingling, headache, sickness, flickering). They rated each side effect in terms of intensity, and likelihood of being related to stimulation on a 5-point Likert scale ranging from 1 – absent to 5 – very strong.

### 2.6 Data analysis

Log files obtained from E-Prime were converted into a single table containing all trials of all participants using Matlab 2025b (The MathWorks Inc, Natick, MA, USA). All subsequent statistical analyses were performed in R 4.5.2 (The R Core Team, R Foundation for Statistical Computing, Vienna, Austria) running in R-Studio 2026.01.0 (Posit Software, MA, USA).

In the pre-registration, we originally planned to analyze DRB effects in RTs and performance. However, due to strong ceiling effects in participants performance (100% accuracy in the majority of the blocks), data analysis focused on RTs in probe trials. Only probe trials with correct responses and correct responses in preceding prime trials were considered. In addition, we excluded prime-probe combinations with prime or probe RTs slower than 1000-ms or faster than 200-ms. Binding scores were computed by averaging reaction times to probe stimuli within experimental conditions (with the factors of STIMULATION, BLOCK, and RSI) as a function of response relation and distractor relation. Average RTs were then used to first calculate the distractor-related effect separately for RC and RR trials and then to add the resulting difference scores:

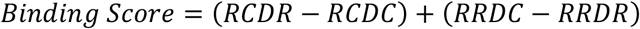

To test for general binding effects before stimulation, and to verify that there were no baseline differences between stimulation sessions, binding scores before stimulation were submitted to a linear mixed-effects model (LLM) using the *lme4* package in R. P-values were obtained using Satterthwaite approximation implemented in the *anova* function of the *lmerTest* package in R, partial eta-squared (η^2^_p_) was obtained from the model as a measure of effect size using the *effectsize* package. The model was fitted with the factors STIMULATION (verum vs. shunt), BLOCK (Block 1 vs. Block 2), RSI (500 vs. 2000) and their respective interactions. In addition, we compared binding scores in the RSI 500 and 2000 conditions in blocks 1 and 2 against zero to test for significant binding in each condition using one sample t-tests (Bonferroni-Holm corrected for multiple comparisons).

To assess tACS effects on binding *during* stimulation, we first subtracted baseline scores in block 2 from scores in subsequent blocks to obtain a score reflecting the change in binding (Δbinding). Change scores were then submitted to an LMM with the same factors as before, except that the factor block, now focused on blocks 3 and 4 (during stimulation). To assess *aftereffects* of tACS, we repeated the same analysis for blocks 5 and 6 (after stimulation). We chose this approach over the originally pre-registered analysis as the normalization relative to pre-stimulation baseline is better suited to handle a large number of post-stimulation blocks, as tACS induced changes in normalized data reflect in a main effect of STIMULATION, rather than a STIMULATION*BLOCK interaction. In addition, normalization reduces the influence of between-session and inter-individual differences in RTs. The use of LMMs instead of ANOVAs further increases sensitivity to effects.

Frequency of sensations and side effects were analyzed by counting the number of intensity ratings > 1 and submitting them to McNemar’s chi-squared test for count data. Intensities of sensations were compared by additionally submitting intensity ratings to Wilcoxon sign-ranked tests.

## 3 Results

### 3.1 DRB effects build up during second half of pre-stimulation baseline

Linear mixed-effects model analysis on baseline binding scores revealed a significant main effect of BLOCK (F_1,301_ = 6.76, p = .010, η^2^_p_ = .02) as well as a significant BLOCK x RSI interaction (F_1,301_ = 5.44, p = .020, η^2^_p_ = .02). One-sided, one-sample t-tests against zero revealed a DRB effect significantly different from zero only in the RSI 500 condition in block 2 (t_43_ = 2.49, p = .033, d = .37, Bonferroni-Holm corrected), but not in block 1 (t_43_ = -2.16, p = .982, uncorrected) or the RSI 2000 condition (all p > .846, uncorrected). While the results reflect the expected modulation of DRB effects under different RSIs, they also suggest a potential build-up of binding effects in the RSI 500 condition from block 1 to block 2 (**Fig. 3a**). Importantly, there were no significant main effects or interactions related to STIMULATION (all p > .21) confirming that there were no significant differences in binding scores before the tACS and shunt applications.

**Figure 3:**
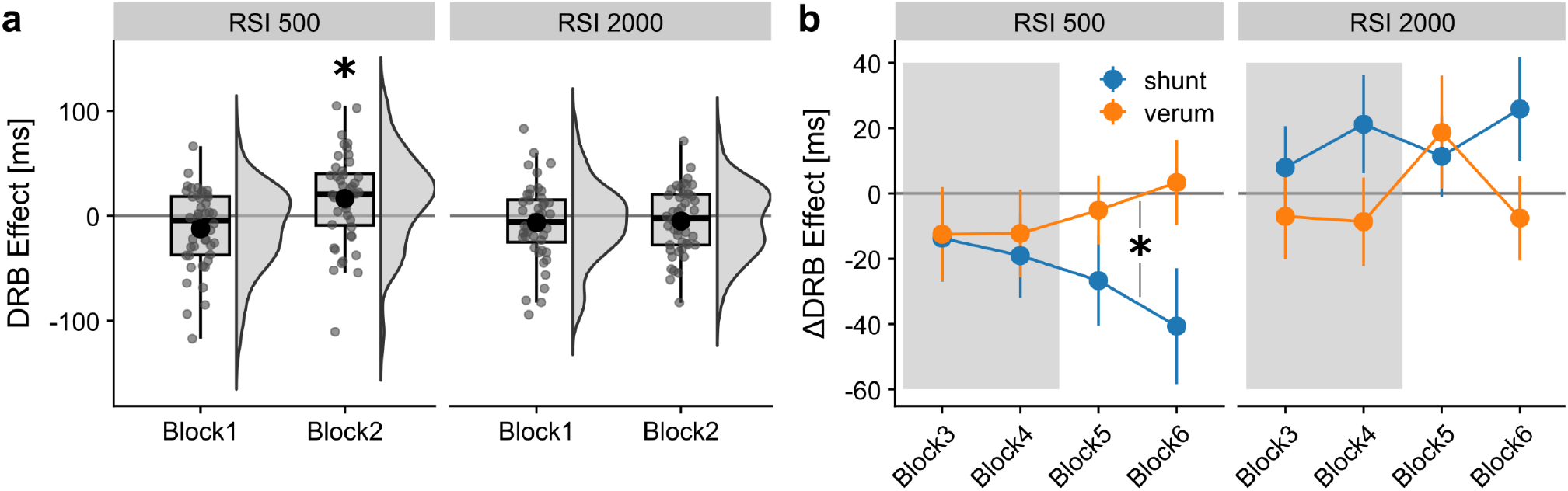
Differential aftereffect of β-tACS on binding scores. **(a)** Average DRB effect during baseline blocks (before tACS or shunt) for RSI 500 and RSI 2000 conditions. Raincloud plots depict individual average binding scores along with boxplots indicating median, 25^th^, and 75^th^ percentile and data range within 1.5 x interquartile range. Asterisk indicates significant deviation of DRB effect from 0. **(b)** Binding score change during (gray area) and after stimulation (white area). Values represent average binding score change relative to block 2. Error bars depict standard error of the mean (S.E.M.). Asterisks indicate significant difference in DRB change after stimulation (main effect in blocks 5 and 6).

### 3.2 No-online effects of β-tACS on DRB

To test for modulation of binding scores *during* stimulation, we performed separate linear mixed-effects model analyses on the change of binding scores relative to block 2 (which turned out to be a sensitive time window for the expected binding effect for the 500-ms RSI condition in the pre-stimulation baseline analysis) for block 3 & 4 and blocks 5 & 6. The analysis of online effects revealed no significant main effects or interactions. Only the main effect of RSI approached significance (F_1, 301_ = 3.64, p = .057, η^2^_p_ = .01). Importantly, the analysis did not reveal any main effect or interaction involving the factor STIMULATION (all p > .156), indicating no evidence for an *online* effect of β-tACS on DRB. Aftereffects of β-tACS on DRB

The analysis of *aftereffects* of β-tACS on binding scores in block 5 and 6 yielded a significant main effect of RSI (F_1, 301_ = 9.24, p_1, 129_ = .003, η^2^_p_ = .03) as well as a significant RSI*STIMULATION interaction (F_1, 301_ = 5.62, p = .018, η^2^_p_ = .02). None of the other main effects or interactions approached significance (all p > .103). To further explore the interaction, we repeated the analysis separately for RSI 500 and RSI 2000 conditions. A significant main effect of STIMULATION was only found in the RSI 500 (F_1, 129_ = 6.58, p = .011, η^2^ = .05) but not in the RSI 2000 condition (F = 0.87, p = .352, **Fig. 3b**).

### 3.3 Perceptual similarity of active vs. shunt tACS

A majority of participants reported mild to moderate cutaneous sensations (itching, tingling, burning) during the experiment. Wilcoxon signed-rank tests for paired samples did not indicate any differences in the intensity rating of sensations between verum and shunt stimulation (all p>.14, uncorrected, **Table 1**). In addition, McNemar’s Chi-squared tests for count data did not indicate any differences in the probability of reporting sensations or side effects between verum and shunt stimulation (all p>.18, uncorrected, **Table 1**), verifying the ability of the shunt montage to induce sensations comparable to active stimulation.

**Table 1.** Overview of sensation and side effect ratings after verum and shunt tACS.

|  | verum |  | shunt |  | p-value | p-value |
| --- | --- | --- | --- | --- | --- | --- |
|  | proportion | intensity | proportion | intensity | intensity | prop. |
| **sickness** | 9.1% | 1.1 ± 0.38 | 13.6% | 1.2 ± 0.54 | .59 | .68 |
| **itching** | 52.3% | 2.0 ± 1.18 | 61.4% | 2.1 ± 1.12 | .49 | .34 |
| **tingling** | 95.5% | 3.0 ± 0.93 | 93.2% | 3.0 ± 1.00 | 1 | 1 |
| **headache** | 27.3% | 1.4 ± 0.69 | 15.9% | 1.2 ± 0.60 | .14 | .18 |
| **burning** | 50.0% | 1.7 ± 0.75 | 47.7% | 1.6 ± 0.72 | .71 | 1 |
| **flickering** | 34.1% | 1.6 ± 1.04 | 34.1% | 1.6 ± 0.95 | .55 | 1 |

*Note:* Shown are proportion of ratings > 1 (i.e., non-absent sensation), average intensity ± SD and statistical comparisons between verum and shunt. All p-values shown are uncorrected.

## 4 Discussion

The current study aimed to investigate the causal role of β-oscillations for the maintenance of perception-action representations (i.e., event files) by modulating β-oscillatory activity with tACS during a DRB task. Contrary to our initial hypothesis, we did not observe stimulation-induced changes in behavioral DRB effects *during* the stimulation period. However, we found a positive effect of β-tACS on DRB *after* stimulation. Importantly, this effect depended on the interval between the prime response and the probe stimulus (RSI), a manipulation commonly used to assess event-file maintenance [10–12]. Replicating previous research, behavioral DRB effects were present at the 500-ms RSI, but not at the 2000-ms RSI [6,12]. Novel to the present study is the finding that, in the sham condition, the DRB effect in the 500-ms RSI condition initially increased (from block 1 to block 2), but then gradually decreased (across blocks 3 to 6). In contrast, in the β-tACS condition, the DRB effect remained elevated during and after stimulation and, by block 6 in the post-stimulation phase, was approximately as large as in pre-stimulation block 2. Together, these findings extend the previous evidence linking β-oscillations to the maintenance of perception-action bindings [12,13] by suggesting a causal contribution of β-oscillations to the stabilization of event files over time. More broadly, the results are consistent with the idea that β-oscillations support the maintenance or re-activation of the current cognitive-motor state, in line with accounts linking β-activity to the preservation of the status quo [33,34].

Effects on behavioral DRB were observed only after stimulation, but not during the stimulation period. Such offline effects are consistent with previous work showing that tACS can induce aftereffects that outlast the stimulation period [21–23]. Aftereffects of tACS are commonly observed, particularly following α-band stimulation, in studies combining tACS with EEG [19,23,29], but have also been reported after β-tACS [22]. It should be noted that different mechanisms have been proposed to underlie online- and aftereffects of tACS. While online effects are usually attributed to neural entrainment [17,18,35,36], aftereffects have been linked to processes of neural plasticity [21–23]. It is possible that online and aftereffects differ not only in their neural basis but also in their consequences for behavior [37]. During stimulation, especially when tACS is applied continuously, externally induced synchronization may interfere with ongoing intrinsic activity especially the dynamic synchronization and desynchronization of β-oscillations. They may thus be less beneficial for behavioral DRB. In addition, participants reported substantial sensations during stimulation, which may have contributed to sensory co-stimulation and distraction [30,32]. By contrast, offline effects may reflect plastic changes that allow intrinsic β-oscillations to be upregulated in a more task- and state-dependent manner [25,38], thereby supporting behavioral DRB more effectively after stimulation has ended. One broader implication of the present findings may concern the relation between binding and learning. Recent theoretical work has proposed that learning may emerge when the strength of a stimulus-response binding exceeds a critical threshold, allowing the initially short-lived trace to become consolidated into longer-term memory [39]. According to this view, the transition from transient binding to more durable learning depends on the interplay of initial binding strength, decay, repetition, and the temporal spacing between repetitions. Against this background, the present results raise the possibility that β-oscillations do not merely support the short-term maintenance of event files but may also help create conditions under which learning becomes more likely. If β-oscillatory activity reflects or supports binding strength [12,13], then β-tACS may have increased the persistence of perception-action bindings and thereby enhanced the likelihood that such bindings remain available across repeated encounters. In this sense, β-tACS may not directly induce learning itself, but rather increase the preconditions for learning by stabilizing event files and reducing their decay over time.

The present findings also speak to the question of anatomical specificity. In the current study, stimulation targeted the left parietal cortex and effects were assessed in the DRB paradigm, in which binding primarily concerns responses and task-irrelevant stimulus features. This fits well with our earlier EEG work showing that maintenance-related β-synchronization over parietal regions is associated with behavioral DRB effects [12]. At the same time, recent evidence suggests that β-oscillations related to perception-action binding are not restricted to parietal sites. Indeed, maintenance-related β-synchronization was also linked to behavioral stimulus-response binding in frontal regions, with source analyses pointing to the supplementary motor area [13]. Importantly, however, the task used in that study differed from the present DRB paradigm in a theoretically relevant way: stimulus-response binding involved not only task-irrelevant features but also repetitions versus alternations of task-relevant stimulus information. One possible interpretation, therefore, is that parietal β-activity is particularly involved in maintaining bindings between responses and task-irrelevant stimulus information, whereas frontal or SMA-related β-activity may play a stronger role in maintaining bindings that include task-relevant stimulus features. Future brain-stimulation studies could test this possibility more directly by comparing parietal and frontal stimulation while systematically contrasting bindings of irrelevant versus relevant stimulus information.

Such an approach may also be clinically informative. Relative to healthy controls, patients with functional movement disorders showed significantly increased behavioral stimulus-response binding together with enhanced maintenance-related post-movement β-synchronization, suggesting a state of perception-action hyperbinding in this disorder [13]. If exaggerated β-synchronization contributes causally to this hyperbinding, interventions aimed at disrupting pathological β-synchronization may help to reduce excessive binding tendencies in these patients. For example, it has been proposed that closed-loop anti-phase tACS can be used to suppress oscillatory activity by destructive interference [40–42]. However, these approaches still face major challenges in estimating oscillatory phase in real-time, and in the presence of concurrent electric stimulation, inducing artifacts that cannot be fully removed from the recorded signals [43–46] — Such approaches may also be relevant for other clinical conditions characterized by abnormal β-synchronization, including Parkinson’s disease [47]. At present, however, this clinical implication remains speculative and requires direct experimental testing.

The present findings should also be considered in light of several limitations. Although the experiment was designed to manipulate event-file maintenance by varying the RSI, the continuous application of β-tACS throughout the stimulation blocks means that it cannot be fully ruled out that the stimulation also affected other action-control processes. For example, β-tACS may have influenced the integration of stimulus-response bindings around prime responses. While binding and retrieval processes are hypothesized to rely primarily on oscillatory activity in other frequency bands, most notably theta and gamma oscillations [7], recent work has also linked alpha/beta oscillations to the integration of event files [48], although these effects were observed in secondary visual cortices that is, in regions located more posteriorly than the targeted stimulation region in the present study. Thus, the present pattern is most consistent with an effect on maintenance in the RSI but does not conclusively exclude an additional contribution of altered integration at the prime. Future studies should capitalize on the behavioral aftereffect observed here and combine tACS with EEG to test more directly whether stimulation-induced changes in β-oscillatory activity are expressed during the RSI and specifically modulate event-file maintenance. A combined tACS-EEG approach may also help to optimize stimulation efficacy by adjusting stimulation frequency to each participant’s individual beta peak. Indeed, previous research indicates that a mismatch between endogenous oscillatory frequency and stimulation frequency may contribute to inter-individual variability in brain-stimulation effects [21,49].

In conclusion, the present study provides causal evidence that β-oscillations contribute to the maintenance of perception-action bindings over time. Although β-tACS did not modulate behavioral DRB during stimulation, it enhanced DRB after stimulation in a manner consistent with reduced event-file decay at the short RSI. These findings extend previous correlational EEG evidence and support the view that β-oscillations help stabilize event files, thereby contributing to the maintenance of perception-action bindings.

## 5 Acknowledgements

We thank Thorsten Brinkmann for assistance with participant acquisition.

## 6 CRediT author statement

**Bernhard Pastötter:** Conceptualization, Methodology, Software, Formal Analysis, Writing – Original Draft, Visualization **Phillipp P. Schummers:** Investigation, Data Curation, Writing – Review & Editing **Paula M. A. Giebel:** Investigation, Data Curation, Formal Analysis, Writing – Review & Editing **Christian Frings:** Conceptualization, Writing – Review & Editing, Funding Acquisition **Florian H. Kasten:** Conceptualization, Methodology, Software, Formal Analysis, Supervision, Investigation, Writing – Original Draft, Visualization

